# Giraffe: a tool for comprehensive processing and visualization of multiple long-read sequencing data

**DOI:** 10.1101/2024.05.10.593289

**Authors:** Xudong Liu, Yanwen Shao, Zhihao Guo, Ying Ni, Xuan Sun, Anskar Yu Hung Leung, Runsheng Li

## Abstract

Third-generation sequencing techniques have become increasingly popular due to their ability to generate long, high-quality reads. Utilizing datasets from various samples and multiple sequencing platforms for comparative and comprehensive analysis is essential for exploring biological mechanisms and establishing benchmark baselines. However, current tools for long reads primarily focus on quality control (QC) and read processing for individual samples, complicating the profiling and comparison of multiple datasets. The lack of tools for data comparison and visualization presents challenges for researchers with limited bioinformatics experience. Furthermore, developing a comprehensive long-read QC method that facilitates comparative analysis and visualization across multiple samples and platforms is necessary to establish benchmark baselines for selecting appropriate sequencing platforms. We introduce Giraffe, a Python3-based command line tool designed for comparative analysis and visualization across multiple samples and platforms. Giraffe enables the assessment of read quality, sequencing bias, and genomic regional methylation proportions for both DNA and direct RNA sequencing reads. Its usability has been demonstrated in various scenarios, including comparisons of different biological processing methods (whole genome amplification vs. shotgun), sequencing platforms (Oxford Nanopore Technology vs. Pacific Biosciences), tissues (kidney marrow with and without blood), and biological replicates (kidney marrows). Additionally, our findings indicate that Oxford Nanopore duplex reads outperform PacBio HiFi reads in homopolymer identification and GC evenness while maintaining comparable overall read quality.

## 1. Introduction

Comparative analysis of samples collected from different individuals or generated from different sequencing platforms is crucial for exploring biological mechanisms and establishing benchmark baselines. Third-generation sequencing platforms, such as Oxford Nanopore Technologies (ONT) and Pacific Biosciences (PacBio), have gained significant popularity in biological research due to their ability to generate long and high-quality reads [1-6]. However, current tools for quality control and long-read assessment, such as pycoQC [7] and minion_qc [8], are primarily designed to analyze in-house sequencing data and handle individual samples, lacking the capability for comparative analysis across multiple samples and platforms.

To address this gap, we developed Giraffe, a set of Python3-based scripts specifically designed for multiple long-read data comparisons in read quality, sequencing bias, and distribution of genomic regional methylation proportion. Compared to alternative tools like NanoComp [9], Giraffe provides greater features and options such as estimated and observed read accuracy, facilitating users in understanding their data quality better and determining whether the quality is reliable enough for downstream analysis.

## 2. Materials and methods

### 2.1 Pre-processing example data

The raw current signal files in fast5 format generated from the ONT platform were profiled to get the base and methylation information using Dorado V0.5.3 (https://github.com/nanoporetech/dorado).

For the read basecalling model, <u>dna_r10.4.1_e8.2_400bps_fast@v4.3.0</u> and <u>dna_r9.4.1_e8_sup@v3.6</u> were used for R10.4.1 and R9.4.1 data, respectively. For the methylation basecalling model, <u>dna_r9.4.1_e8_sup@v3.3_5mCG@v0.1</u> and <u>dna_r10.4.1_e8.2_400bps_sup@v4.3.0_5mC_5hmC@v1</u> were used for R9.4.1 and R10.4.1 data, respectively.

### 2.2 Computing estimating read accuracy

To estimate the read accuracy, we first converted the Q score associated with each base into an error probability, thereby quantifying the likelihood of an error occurring at that specific base position within a read. Then, the average error probability across all bases within a given read was computed and served as a comprehensive representation of the read error probability. To derive the estimated read accuracy, we subtracted the read error probability from 1, as indicated by equation 1.

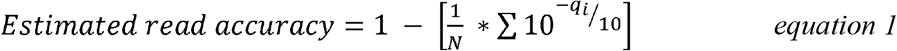

Here, N represents the total number of bases within a read and *q*_*i*_ denotes the base quality score assigned to the *i*-th basecall within the read. This equation was well applied to zebrafish and human samples [6, 10].

### 2.3 Computing observed read quality and mismatch proportion

To get the observed read quality such as accuracy and mismatches proportion. The read was aligned against their reference genome using Minimap2 [11, 12]. The different alignment agreements were used for different sequencing platforms, the “map-ont” and “map-pb” were used for ONT and Pacbio sequencing platforms, respectively. The resulting alignment SAM file was sorted and converted into the BAM file using samtools [13]. The Python package pysam [13-16] was utilized to load the BAM file and take the alignment features. Next, we summarized the length of substitution, insertion, deletion, and matched base within the read and computer the observed accuracy (eq. 3) and mismatches proportion (eq. 5-7).

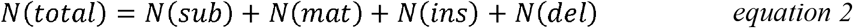

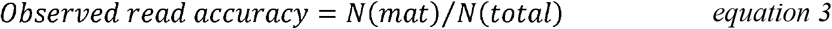

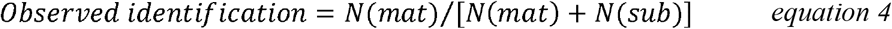

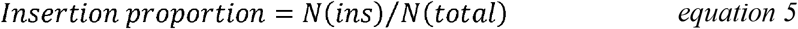

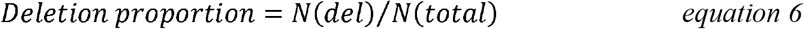

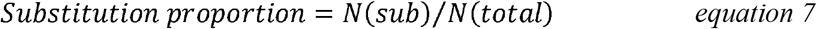

Here *N*(*mat*), *N*(*ins*), *N*(*del*), and *N*(*sub*) were length of matched base, insertion, deletion, and substitution within the read. Those equations were well applied to zebrafish and human samples [6, 10].

### 2.4 Calculating sequencing bias

To detect the sequencing bias within sequencing data, we divide the genome into multiple bins first. The bin size is set to a default value of 1 kb, the user can also provide a different bin size if desired. Within these bins, the “gcbias” function can calculate the GC content and sequencing depth. The bins are then categorized based on their GC content range from 0 to 100%. Then we summarize the distribution of these bins and determine the average read coverage for each GC content range. To ensure effective analysis, the function selects a scale that encompasses over 90% of the bins, as most of the GC content falls within smaller ranges rather than spanning the entire 0 to 100% spectrum. Sequencing depth is subsequently normalized within this selected scale.

## 3. Results and discussion

### 3.1 Software description

Giraffe has four functions, including “estimate” for estimated read quality assessment, “observe” for observed read accuracy evaluation, “gcbias” for sequencing bias detection, and “modbin” for genomic regional methylation proportion comparison, with the requirement of FASTQ reads or alignment SAM/ BAM file (**Fig. 1**).

**Fig. 1.**
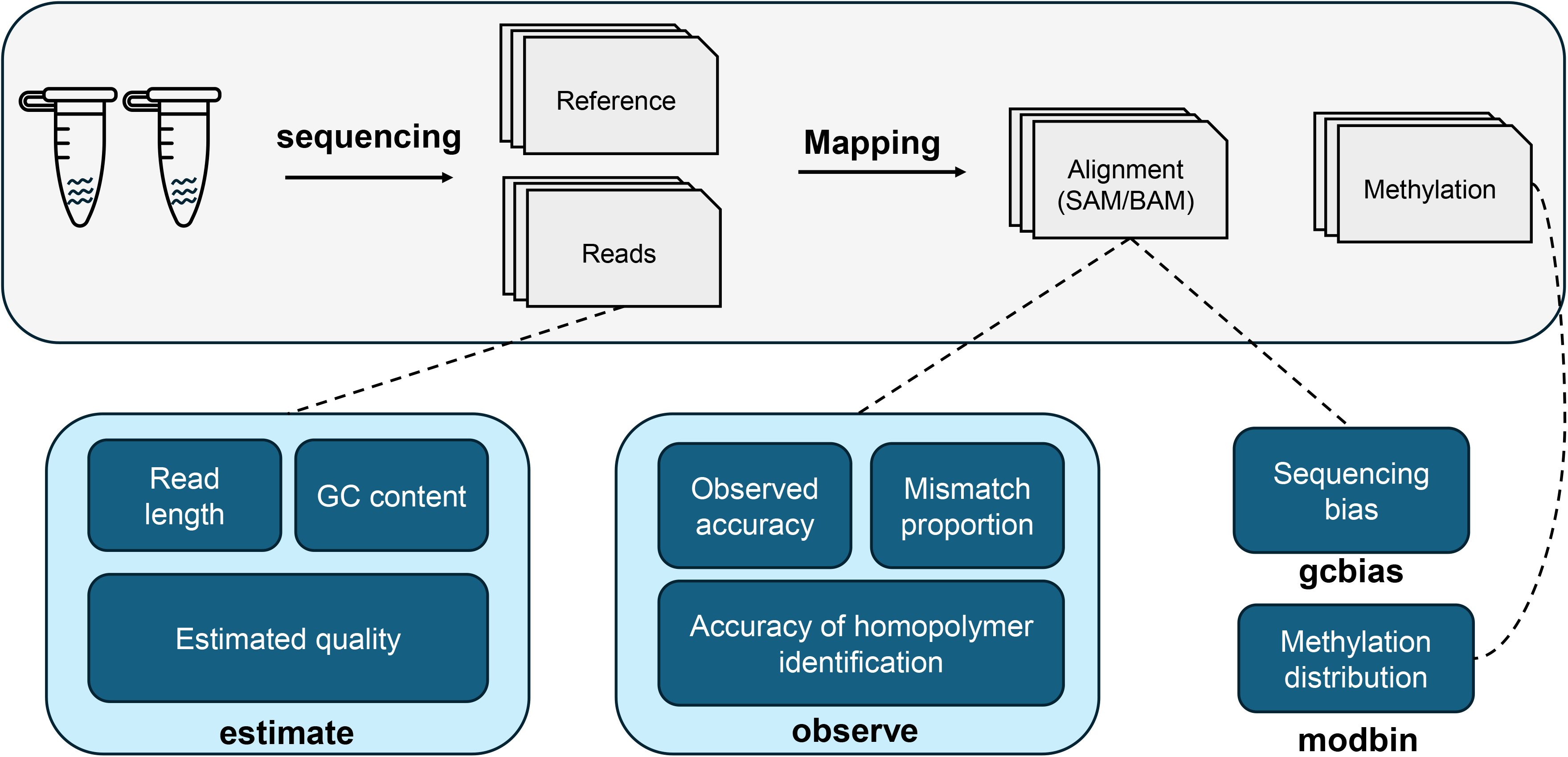
The workflow of the Giraffe. The basic information included in length, GC content, and estimated accuracy for each read can be profiled by the “estimate” function. After aligning reads against the reference, observed accuracy, mismatch proportion, and accuracy of homopolymer identification are calcuslated by the “observe” function. The alignment files can be also used by the “gcbias” function to detect the sequencing bias. As for the genomic regional methylation proportion, it can be summarized by the “modbin” function.

### 3.2 Installation and dependencies

Giraffe and the associated Python scripts are conveniently accessible through the public software repositories PyPI, which can be installed using pip. To support the read processing functionalities, Giraffe relies on dependencies including minimap2 [11, 12], samtools [16], and bedtools [17], which can be obtained through conda (https://conda.org). The scripts are developed utilizing several third-party Python modules, including pysam [13-16], plotnine (https://github.com/has2k1/plotnine), numpy [18], pandas (https://github.com/pandas-dev/pandas), tqdm [19], Biopython (https://biopython.org), and termcolor (https://github.com/termcolor/termcolor).

### 3.3 Estimated read quality

The “estimate” function provides the user with basic information, generating a table that includes estimated accuracy, estimated error proportion, Q score, read length, and GC content (**Table S1**). The function requires input from a table containing sample ID, sequencing platform (ONT/ Pacbio), and the path of the data in FASTQ format. To facilitate simpler and more intuitive comparisons of data quality, we have transformed the Q score into estimated accuracy, providing values ranging from 0 to 100 % (**Equation 1**).

### 3.4 Observe read accuracy

The Q score is calculated by the estimated probability of the base being wrong, which is commonly used to quantify the quality of baseballs in sequencing data. However, there are situations, such as overfitting of the basecalling model, where the Q score may not accurately reflect the read quality, which leads to the development of the “observe” function based on the read alignment (**Equation 2**-**7**). Considering that detecting the homopolymer in long-read sequencing accurately is still challenging, homopolymer identification is also included in the “observe” function as a comparative metric. “Observe” requires a table identical to that of “estimate” and a reference as input, generating two summary tables, one specifically for homopolymer identification accuracy (**Tables S2**) and another for observation information (**Tables S3**) with detailed information such as length of insertions, deletions, substitutions, and matches, as well as the observed read accuracy.

### 3.5 Sequencing bias

In wet lab experiments, some processes can introduce sequencing bias such as polymerase chain reaction (PCR). More specifically, during DNA application, the use of non-random primers or different affinities of polymerase enzyme to DNA sequences can result in uneven coverage and increased depth of specific regions. The results generated by analyzing reads with a sequencing bias are unreliable, especially for methods based on counting. The “gcbias” function was designed to assess the relationship between GC content and sequencing depth, reflecting the level of sequencing bias.

### 3.6 Genomic regional methylation profiling

The methylation proportion in eukaryotes is associated with the strength of promoter activity and downstream protein expression, which has significant implications for exploring the mechanisms underlying various diseases. Nanopore sequencing for native DNA can now give accurate 5mC methylation profiling for every CpG site, especially for human samples. The methylation level for each functional region would be required for downstream analysis. The “modbin” function was designed to facilitate the comparison of methylation profiling files at the genomic regional level and detect the difference between samples.

### 3.7 Additional scripts and options

Considering that the scales of normalization may not be reasonable or suitable for all users in the sequencing bias detection part, additional scripts, including “renormalization_sequencing_bias” and “replot_sequencing_bias”, are provided to allow users to manually define the normalized scale based on the distribution of bins and replot figure. Additionally, another script, named “homopolymer_count.py”, is offered to count the number, position, and type of homopolymers in the reference genome. To support direct RNA sequencing on the ONT platform, we have included the RNA alignment strategy in Giraffe. The user can indicate “ONT_RNA” as the sequencing platform in the input table to assess the quality of RNA data.

### 3.8 Application of Giraffe to different samples

Here, we showcased four applications of Giraffe: exploring the influence of amplification, benchmarking the read quality between different sequencing platforms, identifying the influence of sample purity on sequencing data, and investigating the consistency of biological repetitions. The details of all datasets are available in **Table S4**.

DNA amplification is an essential step before obtaining genome information from limited sample availability. To evaluate the impact of amplification on sequencing reads, DNA samples with and without whole genome amplification derived from the human HCC78 cell line were sequenced using ONT platforms with R9.4.1 flow cells. Upon observation, it became apparent that the amplification process had minimal impact on the sequencing quality and GC content (**Fig. 2A, 2B**, and **2D**-**2F**). However, a noticeable bias was identified in the amplified data (**Fig. 2H**). In terms of methylation proportion at the 1M bin level, a significant majority of the bins derived from amplified reads clustered near 0 percent methylation (**Fig. 2I**).

**Fig. 2.**
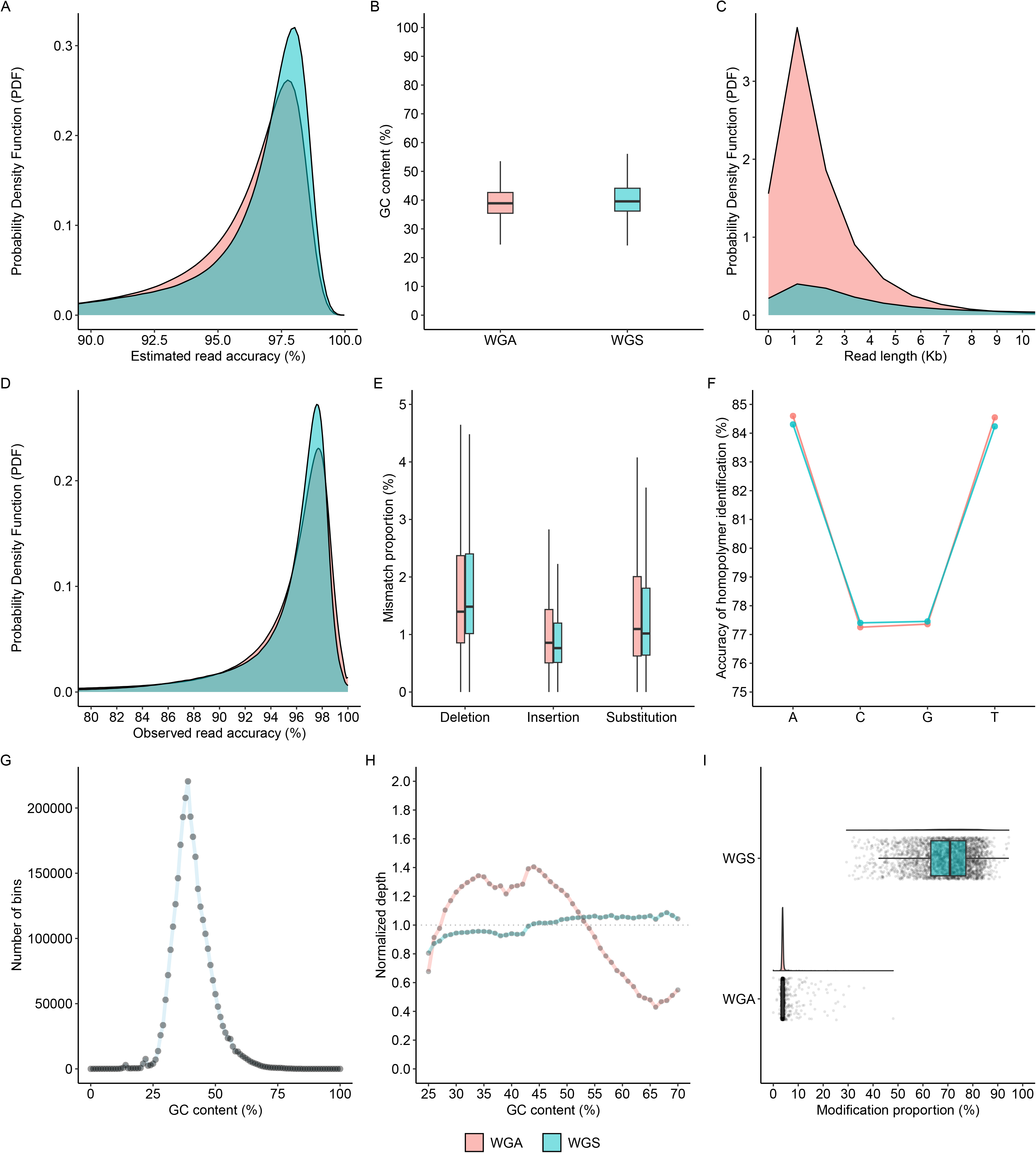
Comparison of human HCC78 cell line between WGS and WGA reads. (**A**), (**B**), and (**C**) the distribution of estimated accuracy, GC content, and length. (**D**) and (**E**) the distribution of observed accuracy and mismatch proportion. (**F**) The accuracy of homopolymer identification for each base type. (**G**) The distribution of 1kp bin numbers within 0 to 100% GC content. (**H**) Relationship between sequencing depth and GC content at 1k bin level. (**I**) The distribution of 5mC methylation proportion at 1M bin level. WGS: whole genome sequencing; WGA: whole-genome amplification.

Different long-read sequencing platforms offer distinct advantages depending on the specific requirements. To assess the read accuracy of long-read sequencing platforms, the ONT duplex reads and PacBio HiFi reads derived from human HG002 datasets were employed. Remarkably, both datasets achieved exceptional accuracy levels exceeding 99% and showcased similar GC content of approximately 40% (**Fig. S1A, S1B**, and **S1D**). Notably, the PacBio reads exhibited an enrichment of around 20 Kb in read length, whereas the ONT reads displayed a more evenly distributed range (**Fig. S1C**). Furthermore, in comparison to HiFi reads, duplex reads demonstrated superior performance in homopolymer identification and less bias in terms of read coverage across the reference genome (**Fig. S1F** and **S1H**).

Different tissues within a biological organism generally exhibit distinct patterns of protein expression and methylation proportion due to their specialized functions. However, isolating a specific tissue without any contamination from other tissues can be challenging in practice. To investigate the impact of other tissue contamination on downstream analysis results, the ONT R9.4.1 reads obtained from the kidney marrow of zebrafish, with and without the presence of blood, were employed. A clear observation was made regarding the decreased quality of kidney marrow with blood (KMB) reads (**Fig. S2A** and **S2D**). The decrease in quality was evidenced by higher mismatch proportion and low homopolymer identification accuracy (**Fig. S2E** and **S2F**).

Biological systems are complex and can be influenced by numerous variables, making it difficult to isolate and control a single changing factor in experiments. The biological repetitions are necessary to evaluate the consistency and reproducibility of their results. Here, two sets of zebrafish kidney marrow duplicate samples sequenced by ONT platforms equipped with R10.4.1 flow cells were used for evaluation. The similar read quality, GC content, and methylation pattern of the two samples can be regarded as important metrics of being a qualified replication (**Fig. S3**).

Considering the large amount of data in the aforementioned datasets, we provided users with selected subsets of datasets to facilitate the operation of Giraffe. These datasets include ONT R10.4.1 and R9.4.1 reads collected from *Escherichia coli* with a reference and 5mC methylation files collected from zebrafish blood and kidney samples. The downloading and running commands are available in the supplementary materials. The size of the demo datasets is approximately 246 Mb, and it takes about 7 minutes to conduct four functions with 4 threads.

## Conclusion

Giraffe is a tool designed for comparing multiple long-read sequencing samples. It offers four main functions: “estimate”, “observe”, “gcbias”, and “modbin” involving read quality, sequencing bias, and genomic regional methylation proportion. Additionally, the practicability and applicability of the Giraffe are well-validated in different research backgrounds.

## Abbreviations

QC: quality control
ONT: Oxford Nanopore Technologies
PacBio: Pacific Biosciences
PCR: polymerase chain reaction
WGS: Whole-genome sequencing
WGA: whole-genome amplification

## Data and materials availability

The source code is available at https://github.com/lrslab/Giraffe_View and PyPI (https://pypi.org/project/Giraffe-View).

All the datasets used in this work can be found in **Table S4**. The PacBio HiFi and ONT duplex reads are published, which are available at https://humanpangenome.org/data. The other three datasets can be found at the National Centre for Biotechnology Information (NCBI) with project ID PRJNA1095401. The testing demo datasets are available in Figshare (https://figshare.com/articles/dataset/demo_datasets/25378693).

## Acknowledgments

We thank the High-Performance Computing Cluster at the City University of Hong Kong for providing us with computational resources. We thank Fangfang He and Ruan Yao from The University of Hong Kong for the zebrafish kidney marrow DNA extraction.

## Authors’ contributions

Xudong Liu: Conceptualization, methodology, software, validation, formal analysis, investigation, data curation, writing (original draft, review, and editing). Yanwen Shao: Validation, investigation, data curation, writing (review and editing). Zhihao Guo: Methodology and writing (review and editing). Ying Ni: Data curation, Writing (review and editing). Xuan Sun: Data curation, Writing (review and editing). Anskar Yu Hung Leung: Writing (review and editing), supervision, funding acquisition. Runsheng Li: Conceptualization, methodology, validation, investigation, writing (review, and editing), supervision, and funding acquisition.

## References

[1] Daniel Branton, David W Deamer, Andre Marziali, Hagan Bayley, Steven A Benner et al., The potential and challenges of nanopore sequencing, Nature biotechnology, vol. 26, no. 10, (2008), pp. 1146–1153.

[2] Yunhao Wang, Yue Zhao, Audrey Bollas, Yuru Wang, and Kin Fai Au, Nanopore sequencing technology, bioinformatics and applications, Nature biotechnology, vol. 39, no. 11, (2021), pp. 1348–1365.

[3] Anthony Rhoads, and Kin Fai Au, PacBio Sequencing and its Applications, Genomics, Proteomics & Bioinformatics, vol. 13, no. 5, (2015), pp. 278–289.

[4] Søren M Karst, Ryan M Ziels, Rasmus H Kirkegaard, Emil A Sørensen, Daniel McDonald et al., High-accuracy long-read amplicon sequences using unique molecular identifiers with Nanopore or PacBio sequencing, Nature methods, vol. 18, no. 2, (2021), pp. 165–169.

[5] Peng Ni, Fan Nie, Zeyu Zhong, Jinrui Xu, Neng Huang et al., DNA 5-methylcytosine detection and methylation phasing using PacBio circular consensus sequencing, Nature communications, vol. 14, no. 1, (2023), pp. 4054.

[6] Xudong Liu, Ying Ni, Dandan Wang, Silin Ye, Mengsu Yang et al., Unraveling the whole genome DNA methylation profile of zebrafish kidney marrow by Oxford Nanopore sequencing, Scientific data, vol. 10, no. 1, (2023), pp. 532.

[7] Adrien Leger, and Tommaso Leonardi, pycoQC, interactive quality control for Oxford Nanopore Sequencing, Journal of Open Source Software, vol. 4, no. 34, (2019), pp. 1236.

[8] Robert Lanfear, Miriam Schalamun, David Kainer, Weiwen Wang, and Benjamin Schwessinger, MinIONQC: fast and simple quality control for MinION sequencing data, Bioinformatics, vol. 35, no. 3, (2019), pp. 523–525.

[9] Wouter De Coster, and Rosa Rademakers, NanoPack2: population-scale evaluation of long-read sequencing data, Bioinformatics, vol. 39, no. 5, (2023).

[10] Ying Ni, Xudong Liu, Zemenu Mengistie Simeneh, Mengsu Yang, and Runsheng Li, Benchmarking of Nanopore R10. 4 and R9. 4.1 flow cells in single-cell whole-genome amplification and whole-genome shotgun sequencing, Computational and Structural Biotechnology Journal, vol. 21, (2023), pp. 2352–2364.

[11] H. Li, Minimap2: pairwise alignment for nucleotide sequences, Bioinformatics, vol. 34, no. 18, Sep 15, (2018), pp. 3094–3100.

[12] H. Li, New strategies to improve minimap2 alignment accuracy, Bioinformatics, vol. 37, no. 23, Dec 7, (2021), pp. 4572–4574.

[13] P. Danecek, J. K. Bonfield, J. Liddle, J. Marshall, V. Ohan et al., Twelve years of SAMtools and BCFtools, Gigascience, vol. 10, no. 2, Feb 16, (2021).

[14] pysam-developers, pysam, vol. https://github.com/pysam-developers/pysam.

[15] J. K. Bonfield, J. Marshall, P. Danecek, H. Li, V. Ohan et al., HTSlib: C library for reading/writing high-throughput sequencing data, Gigascience, vol. 10, no. 2, Feb 16, (2021).

[16] H. Li, B. Handsaker, A. Wysoker, T. Fennell, J. Ruan et al., The Sequence Alignment/Map format and SAMtools, Bioinformatics, vol. 25, no. 16, Aug 15, (2009), pp. 2078–9.

[17] A. R. Quinlan, and I. M. Hall, BEDTools: a flexible suite of utilities for comparing genomic features, Bioinformatics, vol. 26, no. 6, Mar 15, (2010), pp. 841–2.

[18] C. R. Harris, K. J. Millman, S. J. van der Walt, R. Gommers, P. Virtanen et al., Array programming with NumPy, Nature, vol. 585, no. 7825, Sep, (2020), pp. 357–362.

[19] Casper O. da Costa-Luis, tqdm: A fast, extensible progress meter for python and cli, Journal of Open Source Software, vol. 4, no. 37, (2019), pp. 1277 %@ 2475-9066.

