## Supplementary material for "Giraffe: a tool for comprehensive processing and visualization of multiple long-read sequencing data": Fig. S1, S2, and S3

#### Supplementary commands

The commands for demo downloading.

```
# The input file list.
wget https://figshare.com/ndownloader/files/44967445 -O fastq.list
wget https://figshare.com/ndownloader/files/44967442 -O bed.list
wget https://figshare.com/ndownloader/files/44967499 -O bam.list

# The reference and ONT reads (R10.4.1 and R9.4.1) of E. coli.
wget https://figshare.com/ndownloader/files/44967436 -O Read.tar.gz

# The 5mC methylation files of zebrafish blood and kidney samples.
# The position file only includes the gene promoter region in chromosome 1.
wget https://figshare.com/ndownloader/files/44967427 -O Methylation.tar.gz

tar -xzvf Read.tar.gz
tar -xzvf Methylation.tar.gz
rm Read.tar.gz Methylation.tar.gz
```

The commands for the Giraffe running.

```
giraffe estimate --input fastq.list --plot --cpu 4
giraffe observe --input fastq.list --plot --cpu 4 --ref Read/ecoli_chrom.fa
giraffe gcbias --input bam.list --plot --ref Read/ecoli_chrom.fa
giraffe modbin --input bed.list --cpu 4 --plot --pos Methylation/zf_promoter.db
```

### Supplementary Figures

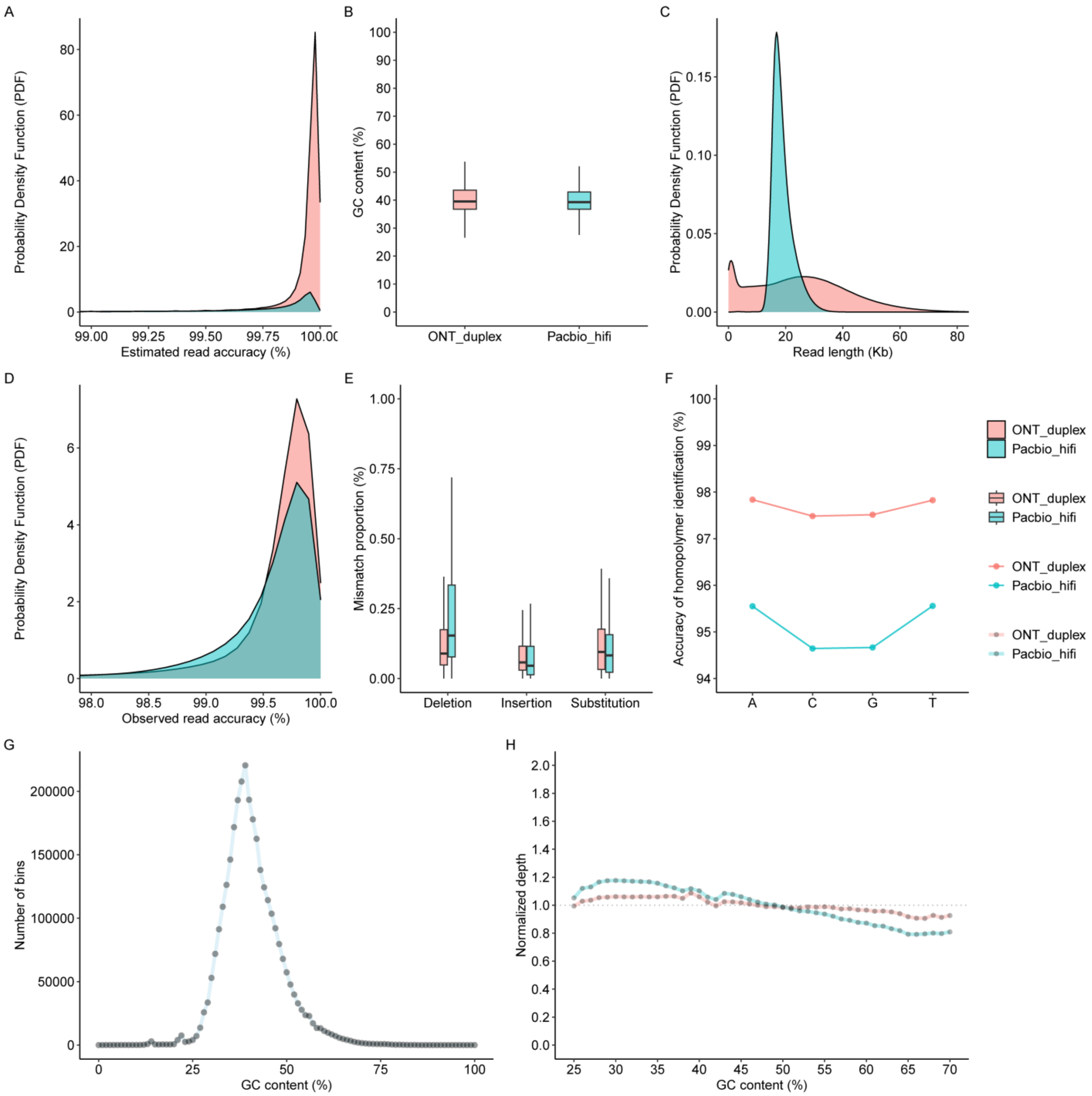


**Figure S1.** Comparison of human HG002 PacBio HiFi and ONT duplex data. (**A**), (**B**), and (**C**) the distribution of estimated accuracy, GC content, and length for each read, respectively. (**D**) and (**E**) the distribution of observed accuracy and mismatch proportion. (**F**) The accuracy of homopolymer identification for each base type. (**G**) The distribution of 1kp bin numbers within 0 to 100% GC content. (**H**) Relationship between sequencing depth and GC content at 1k bin level.

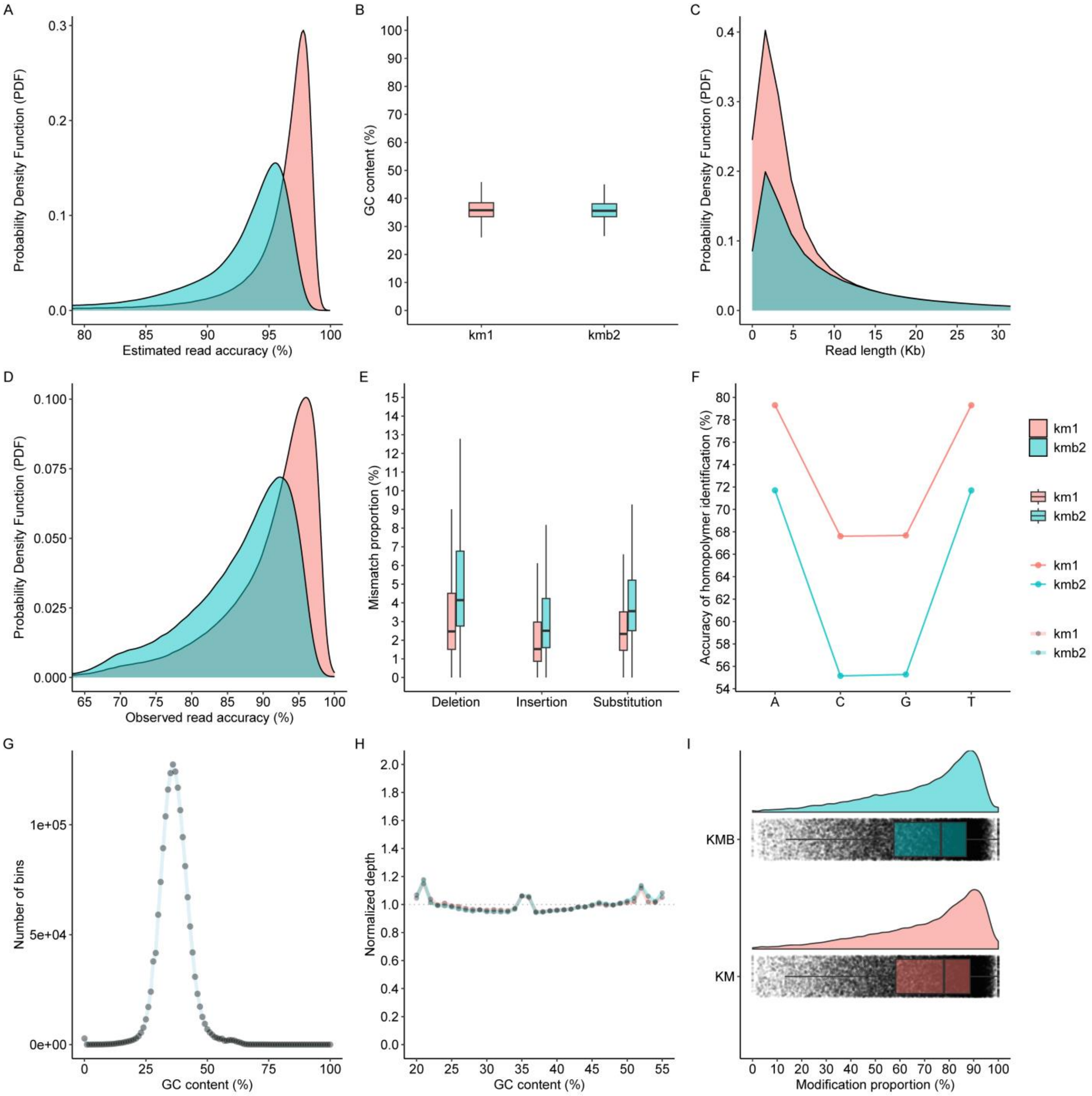


**Figure S2.** Comparison of zebrafish kidney marrow data, with or without blood. (**A**), (**B**), and (**C**) the distribution of estimated accuracy, GC content, and length for each read, respectively. (**D**) and (**E**) the distribution of observed accuracy and mismatch proportion. (**F**) The accuracy of homopolymer identification for each base type. (**G**) The distribution of 1kp bin numbers within 0 to 100% GC content. (**H**) Relationship between sequencing depth and GC content at 1k bin level. (**I**) The distribution of methylation proportion at the promoter level. **km:** kidney marrow, **kmb:** kidney marrow with blood.

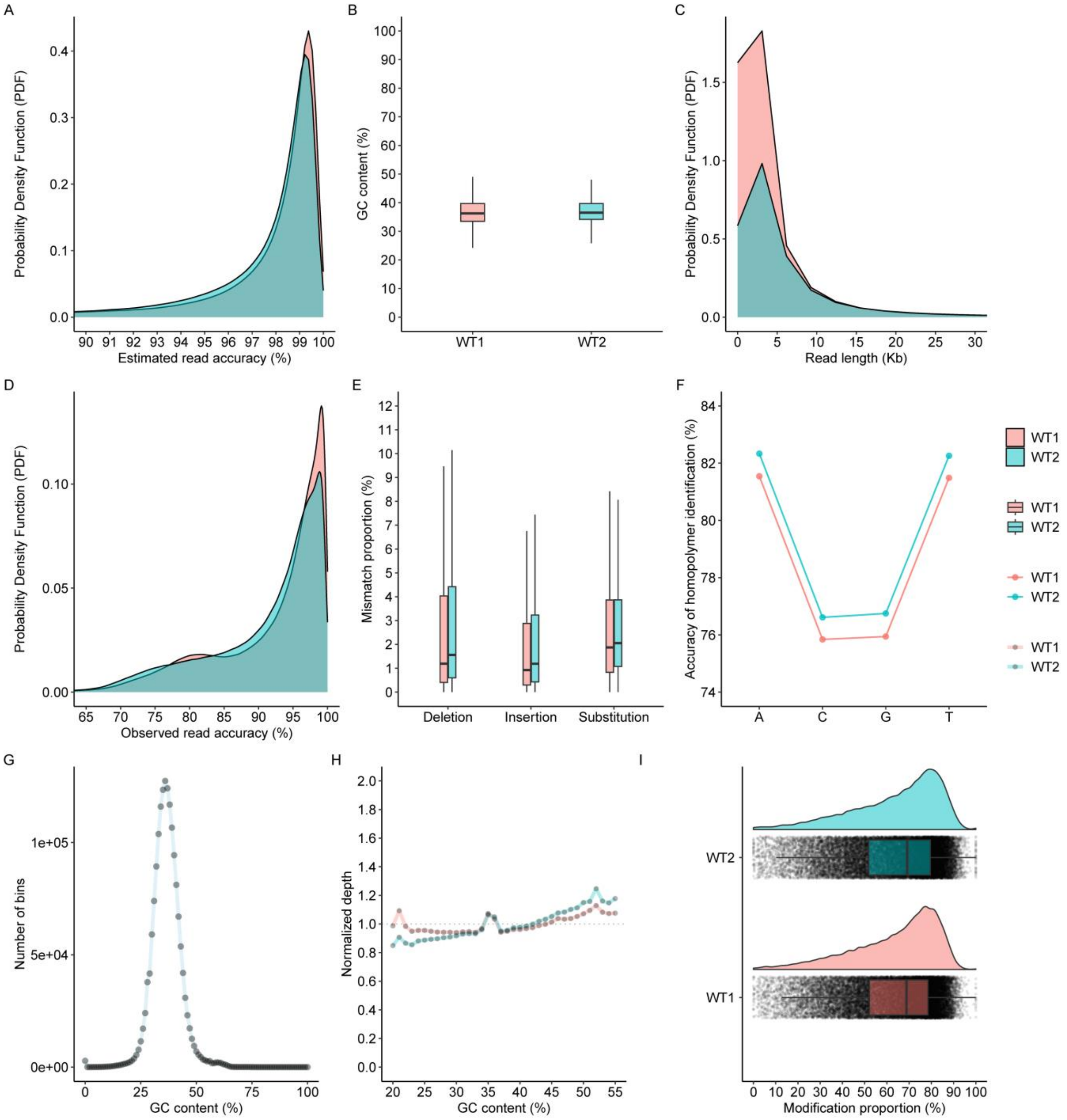


**Figure S3.** Comparison of two zebrafish kidney marrow data. Both samples were collected from the wild type. (**A**), (**B**), and (**C**) the distribution of estimated accuracy, GC content, and length for each read, respectively. (**D**) and (**E**) the distribution of observed accuracy and mismatch proportion. (**F**) The accuracy of homopolymer identification for each base type. (**G**) The distribution of 1kp bin numbers within 0 to 100% GC content. (**H**) Relationship between sequencing depth and GC content at 1k bin level. (**I**) The distribution of methylation proportion at the promoter level.
